# Data-mining of Antibiotic Resistance Genes Provides Insight into the Community Structure of Ocean Microbiome

**DOI:** 10.1101/246033

**Authors:** Shiguang Hao, Pengshuo Yang, Maozhen Han, Junjie Xu, Shaojun Yu, Chaoyun Chen, Wei-Hua Chen, Houjin Zhang, Kang Ning

**Affiliations:** Key Laboratory of Molecular Biophysics of the Ministry of Education, College of Life Science and Technology, Huazhong University of Science and Technology, Wuhan, Hubei, 430074, China

**Keywords:** data-mining, marine microbiome, antibiotic resistance gene, human impact

## Abstract

**Background:** Antibiotics have been spread widely in environments, asserting profound effects on environmental microbes as well as antibiotic resistance genes (ARGs) within these microbes. Therefore, investigating the associations between ARGs and bacterial communities become an important issue for environment protection. Ocean microbiomes are potentially large ARG reservoirs, but the marine ARG distribution and its associations with bacterial communities remain unclear.

**Methods:** we have utilized the big-data mining techniques on ocean microbiome data to analysis the marine ARGs and bacterial distribution on a global scale, and applied comprehensive statistical analysis to unveil the associations between ARG contents, ocean microbial community structures, and environmental factors by reanalyzing 132 metagenomic samples from the *Tara* Oceans project.

**Results:** We identified in total 1,926 unique ARGs and found that: firstly, ARGs are more abundant and diverse in the mesopelagic zone than other water layers. Additionally, ARG-enriched genera are closely connected in co-occurrence network. We also found that ARG-enriched genera are often more abundant than their ARG-less neighbors. Furthermore, we found that samples from the Mediterranean that is surrounded by human activities often contain more ARGs.

**Conclusion:** Our research for investigating the marine ARG distribution and revealing the association between ARG and bacterial communities provide a deeper insight into the marine bacterial communities. We found that ARG-enriched genera were often more abundant than their ARG-less neighbors in the same environment, indicating that genera enriched with ARGs might possess an advantage over others in the competition for survival in the oceanic microbial communities.

## Background

Marine microbial communities represent one of the most abundant and complex communities on earth. Many studies on microbial communities of surface ocean waters [1, 2] have revealed a large reservoir of genes and functional modules [3]. These rich resources have been used for deep data mining [4, 5]. For example, by comparing the metagenomic data qualitatively and quantitatively, fluctuations in taxonomical composition and metabolic capabilities from various environments could be revealed [6]. In consideration of this valuable information, further investigations in complex integral biochemical metabolic processes reflecting the ways in which microbes are accustomed to changing environments should be collated and reported.

The *Tara* Oceans project is so far one of the largest expeditions to collect marine samples [7]. Over the past few years, this project has collected over 30,000 samples from more than 200 sampling sites [8], more than 500 high quality samples have been sequenced by whole genome sequencing (WGS) [9]. These resources provide scientists with valuable information for exploring metabolic pathways involved in biogeochemical cycles at the sampling sites and revealing complex interplays within the microbial communities and between the communities as a whole and the surrounding environments [10].

Ocean microbiomes are potentially large pools of antibiotics and antibiotic resistance genes (ARGs) [11]. ARGs are important to protect bacteria from antibiotics produced by other bacteria and other organisms, and is a key determinant to the dynamic balance of the bacterial community [12, 13]. Antibiotics have been widely used not only in bacterial infection treatment, but also in agriculture and animal husbandry for quite some time [13]. Our research for investigating the marine ARG distribution and revealing the association between they 1) alter the community structure by killing some species that have no resistance to them [14]; other changes may follow because of complex interplays among species, and 2) promote the exchange of ARGs among species [15, 16], which might in turn alter the community structure. Long-term impacts include faster evolution of ARGs [17, 18] and the rise of multidrug-resistance bacteria. Therefore research on antibiotic and ARGs have become more and more important worldwide [19, 20]. How to utilize antibiotics and control antibiotic resistance has become an increasingly important issue [21, 22], especially at industrial settings [23, 24].

Mechanisms of resistance to antibiotics in bacteria have only been revealed recently, thanks to the isolation and genetic characterization of bacteria with ARGs [25]. Many experimental and bioinformatics methods for identifying new antibiotics and ARGs have been developed [26, 27]. Further understanding of the functions of ARG products and their effects on the bacterial community may uncover new ways of the influence of antibiotics and ARGs on natural bacterial communities [16]. However, without advanced data-mining techniques, current studies on identification and annotation of ARG from ocean microbiome data remain illusive.

In this study, in order to reveal the associations between microbiota community structures and ARGs, we have utilized data-mining techniques to reanalyze 132 metagenomic samples from the *Tara* Oceans project, and examined the taxonomical structures as well as functional profiles. The enrichment of ARGs in several marine genera was investigated. Firstly, we identified in total 1,926 unique ARGs and found that the ARG contents were strongly associated with the depth: ARGs were more abundant and diverse in the mesopelagic zone than other water layers. Secondly, ARG-enriched genera, including *Flavobacterium, Alteromonas, Pseudoalteromonas* were closely connected in co-occurrence network and are biomarkers of their respective environments. Thirdly, ARG-enriched genera, such as *Alteromonas, Pseudoalteromonas, Marinobacter*, and *Flavobacterium*, were often more abundant than their ARG-less neighbors. Finally, the relationship between taxonomical structures and ARGs was exemplified in *Flavobacterium*, a common marine genus which was identified as a hub node in species-species co-occurrence network. We detected the enrichment of a resistance type (*bac*A) against bacitracin in *Flavobacterium* using computational approaches and validated the results using statistical tests. Inspired by this example, we attempted to interpret how ARG enrichment occurred in many organisms and thus affected the bacterial community structure, and we hypothesized the significance of human involvement in this, and densely populated Mediterranean was exemplified to prove the ARG effect on bacterial community structure.

## Results and Discussions

### Taxonomical analysis revealed key determinants of community compositions

To facilitate the identification of ARGs and the comparison of ARG contents within and between communities (i.e. samples), we first identified the community compositions (i.e. the number of species and relative abundance of each species) for all the oceanic samples we obtained from the *Tara* Ocean project, and characterized the correlations between community structure and environmental factors, as well as between community structure and species co-occurrence patterns.

#### Microbial community composition and function analysis

We obtained in total 36, 356 microbial OTUs including 715 archaeal and 35,641 bacterial OTUs, respectively. Microbial community profiles at phylum and genus level were illustrated in **Supplementary Fig. S1**. We identified in total 15 phyla and 24 genera that were relative abundant, i.e. with relative abundance above 0.1% (for details please check **Supplementary Table S1)**. Functional analyses on specified KEGG pathway [28] level 2 and level 3 were illustrated in **Supplementary Fig. S2**

#### Species co-occurrence network analysis

To better understand the interactions and associations within the microbial communities, we constructed species co-occurrence networks at genus and OTU level (**Fig. 1a and 1c**). We obtained a network at the genus level (Pearson threshold ±0.1) consisting 20 nodes and 130 edges, with a clustering coefficient of 0.744 and a network density of 0.684. With depth-related information and their first neighbor in network on genus level, a sub-network (**Fig. 1b**) with 11 nodes (6 in surface water layer and 5 in mesopelagic zone) and 52 related edges was selected to exemplify the validity of the network (**Fig. 1a**). The 6 surface nodes and the 5 mesopelagic nodes had strong negative correlations, but in contrast, the nodes within surface water layer or mesopelagic zone showed strong positive correlations. These differences are reasonable, as symbiosis plays a leading role in the same environment, yet such symbiosis patterns might differ greatly in different environments [29]. On OTU level, a connected network with 130 nodes and 3,101 edges was constructed, which had a clustering coefficient of 0.63 and a network density of 0.3 (**Fig. 1c**). The largest cluster colored in black was mainly composed of species from phylum *Proteobacteria*, which was the most abundant phylum in the ocean [10]. We identified four hub nodes in this network, among which two were unclassified species of genera *Flavobacterium* and *Polaribacter* and the other two belonged to phylum *Proteobacteria*.

**Figure 1.**
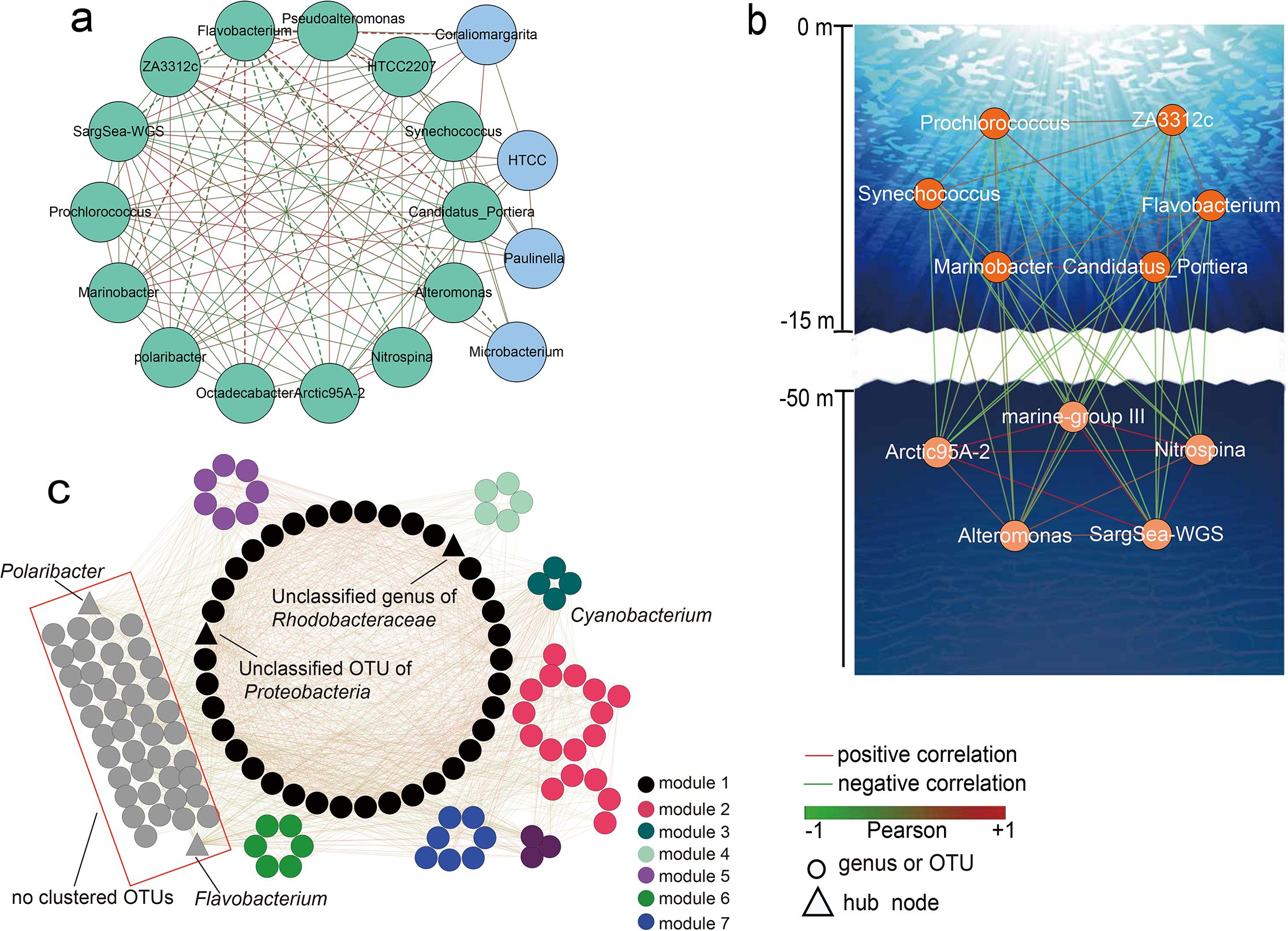
Global views at genus level and OTU level and a subnetwork at genus level. **(a)** The global species co-occurrence network at genus level. Red and green edges represent positive and negative correlation between two linked genera (nodes), respectively. Genera in a cluster were colored in green, while singletons were colored in blue. **(b)** A sub-network related to depth variable at genus level. Depth was an important environment factor and had certain correlations with temperature, oxygen and chlorophyll concentration, so this depth-related sub-network was exemplified the validity for our network. Each node represents a genus and each edge presents a co-occurrence relationship. Color of edges present the relationship strength (calculated by Pearson Correlation Coefficient) of species-species (or genus-genus) co-occurrence relationship. The cluster from surface water contained 6 genera that were highly positively related, and the cluster from deep sea contains 5 genera that were highly positively related. **(c)** The global view of species co-occurrence network at OTU level. 7 clusters labeled in different colors were produced by using MCODE cluster algorithm. Each node represents a selected OTU, and edges in red and green represent positive and negative correlation between two connected OTUs, respectively. The four triangle-shaped nodes were identified as hub nodes in the network.

Genus *Flavobacterium* has been identified as a biomarker (depth-and oxygen-related strategies, *p*-value=5.96e-5 and 2.08e-7, respectively) and a hub node in co-occurrence network, the importance of which was confirmed by previous studies: it is strictly aerobic and tended to live in surface water with high-concentration of chlorophyll and phytoplankton [30, 31], and played an important role in carbon cycling in bacterial communities [32].

### Distribution of antibiotic resistance genes across water layers

By searching 81,850,381 protein sequences from 132 samples against the ARDB database [33], 1,926 unique ARGs were detected (**Supplementary Table S2**). These sequences account for only 0.024‰ of all predicted proteins, which is much lower than that of the human gut microbiome [27]. The 1,926 unique ARGs were classified into 70 different types according to their gene names. This resulted in 27 multidrug types (efflux-mediated), 38 single-drug types (non-efflux), and 5 target-specific types (efflux-mediated). Of the 132 samples, 126 (95.4%) contain at least one ARG sequence (**Supplementary Table S3**).

We correlated the ARG-contents with water layers in order to investigate how ARG distribution was affected. The samples were collected from three layers: surface water layer (SRF), deep chlorophyll maximum layer and subsurface epipelagic mixed layer (DCM/MIX), and mesopelagic zone (MES). We found that among three water layers, SRF and DCM/MIX harbored 44 and 39 resistance types, respectively, while MES harbored 59 resistance types (**Supplementary Table S4**), suggesting there were more resistance types in the deeper water layer. For example, dataset ERS490633 from MES had 26 resistance types, which was the largest amount in a single dataset, while 11 datasets (9 from SRF, one from DCM/MIX and one from MES) had only one resistance type (**Supplementary Table S3**). To eliminate biases due to sequencing depths, we normalized the number of resistance types and ARG sequences in each dataset by the number of processed reads and the number of OTUs (**Supplementary Table S3, Fig. 2a and 2b**). The results showed that the mean of normalized number of resistance types in MES (0.000991) was significantly higher than that in SRF (0.000297) and DCM/MIX (0.000415), with *p*-value=4.251e-11 and 3.836e-9, respectively (Mann-Whitney test); but the difference between SRF and DCM/MIX was not significant (Mann-Whitney test, *p*-value=0.01429>0.01). The mean of normalized number of ARG sequences in MES (0.002439) was significantly higher than that in SRF (0.000525) and DCM/MIX (0.000875), with *p*-value=1.031e-11 and 8.843e-9, respectively (Mann-Whitney test); and the difference between SRF and DCM/MIX was also significant (*p*-value=2.202e-3). Together, these results suggested that ARGs in MES were significantly more diverse; and the diversity increased when the sampling proceeds to deeper zones. And the increasing species richness was also detected when the sampling proceeds to deeper zones according to our biodiversity statistic and previous research for *Tara* Oceans analysis [10, 34]. With limited carbon source and high mobility of mesopelagic zone, the bacteria had a low growth speed but can escape the predator and viral infect [35].

**Figure 2.**
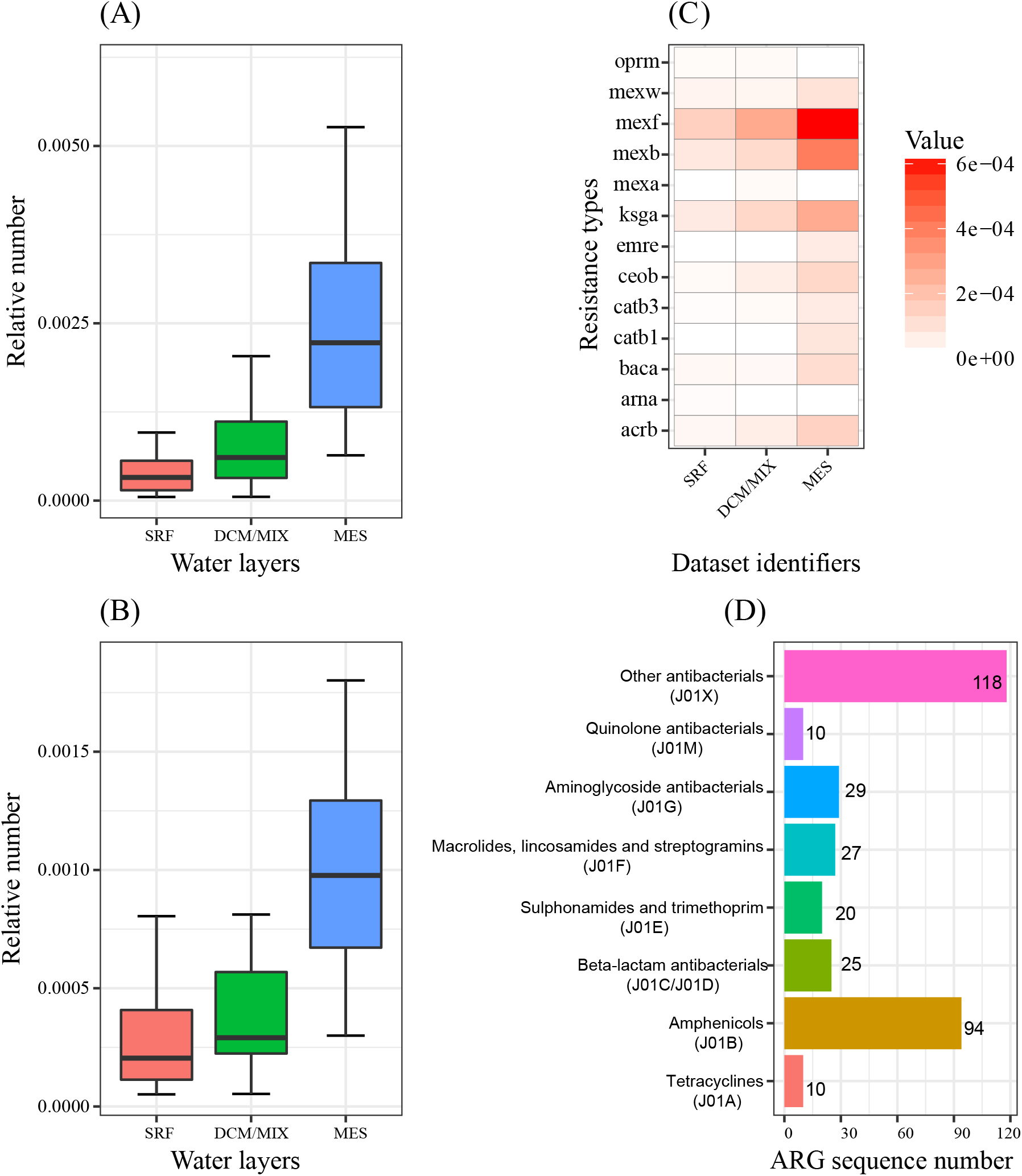
Distribution and classification of detected ARGs. (A) and (B) Boxplots of the distribution of ARG sequences and ARG types in three water layers, respectively. The normalization method was described in section “**Materials and Methods**“. (C) A heatmap of the Top 10 abundant ARG types in each water layer. A white tile means that this ARG type was not detected in this water layer. (D) **The classification of ARGs sequences**. The ARGs sequences are classified according to WHOCC ATC/DDD Index. Amphenicols was the most abundant antibiotic class. Abbreviations used: SRF, surface water layer; DCM, deep chlorophyll maximum layer; MIX, subsurface epipelagic mixed layer; MES, mesopelagic zone. The data used for plotting was exhibited in **Supplementary Table S10**.

The 70 resistance types were unevenly distributed among the three water layers (**Supplementary Fig. S7**). For example, *mex*F was present in 41 out of 55 datasets (74.5%) in SRF, 40 of 42 datasets in DCM/MIX (95.2%), and all 29 datasets in MES (100%) (**Supplementary Table S5**), while 5, 2, and 17 types were found to be specific to SRF, DCM/MIX, and MES, respectively (**Supplementary Table S4**). The top 10 most abundant resistance types in each layer were plotted in **Fig. 2c**. All top 10 resistance types in MES were present in more than half of datasets, while only 2 and 4 of the top 10 resistance types in SRF and DCM/MIX were present in more than half of datasets, respectively (**Supplementary Table S5**). This result indicates the resistance types in MES are distributed more widely. The following multidrug resistance types, including *mex*F, *mex*B, *acr*B, *ceo*B, and *mex*W, were found in the top 10 of three layers, with a high abundance, which suggests that multidrug resistance types are abundant and common and have important contributions to antibiotic resistance [36].

To investigate the antibiotic resistance gene classification, the 1,926 unique ARGs were mapped according to WHOCC ATC/DDD Index (https://www.whocc.no/atc_ddd_index/?code=J01) and the relative abundances of types conferring resistance to the same antibiotic were calculated (**Fig. 2d**). Only 333 of the 1,926 ARG sequences were classified. The excluded sequences are 228 *ksg*A sequences, for which we cannot find a proper Index, and 1,365 multidrug efflux pumps.

### ARG-enriched genera and their connection with biomarkers and co-occurrence network

As a result of taxonomical assignment of ARGs, we successfully assigned 1,659 unique ARGs to 11 genera, which could be classified into 75 resistance types (**Supplementary Table S6)**. The enrichment of ARGs at genus level was exemplified by the 20 resistance types illustrated in **Fig. 3** (see **Supplementary Table S6 and S7** for all the 75 resistance types). To determine whether a resistance type was enriched in a genus, univariate hypergeometric tests (**Fig. 3a**) were applied on each resistance type against each genus, with results showing that ARGs of 37 resistance types were found enriched in at least one genus (*p*-value<0.01). Meanwhile, to determine whether a genus was enriched with ARGs, multivariate hypergeometric tests were applied on all the resistance types against each genus, with results showing that 4 genera were well enriched with ARGs, including *Marinobacter* (*p*-value=6.82e-201), *Alteromonas* (*p*-value=8.28e-198), *Flavobacterium* (*p*-value=5.90e-143), and *Pseudoalteromonas* (*p*-value=3.25e-101) (**Fig. 3d**), and these 4 genera indeed harbored most ARGs (435, 515, 101 and 602 respectively). To determine whether a resistance type is enriched in all genera, multivariate hypergeometric tests (**Fig. 3b**, **Supplementary Table S8**) on each resistance type was performed again, which revealed that *bac*A was the third enriched type in these genera (*p*-value=1.67e-63), behind *mex*F and *ksg*A (*p*-value=3.84e-96 and 3.30e-72, respectively).

**Figure 3.**
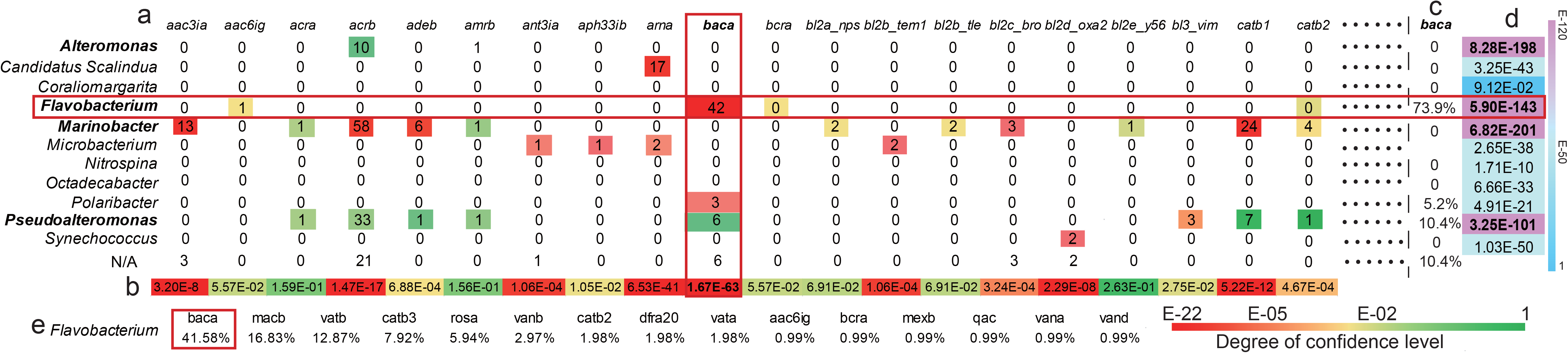
Enrichment analysis of ARGs in marine microbial genera. A total of 20 out of 75 resistance types were selected as examples to show the enrichment of ARGs in genera (the complete set of data used was exhibited in **Supplementary Table S7**). **(a)** To determine whether a resistance type is enriched in a genus, univariate hypergeometric test is performed. The cell color is determined according to the *p*-values produced by univariate hypergeometric tests. Column names represent resistance types and row names represent genera. A “N/A” tag was assigned to a row that contains ARGs that are not identified in any of the 11 genera or the best hit did not meet the identity threshold of 40%. The horizontal and vertical rectangles highlight the number of ARGs in *Flavobacterium* and the number of *bac*A in genera, respectively. In the cell where two rectangles overlap, the number means that 42 *bac*A sequences were identified in *Flavobacterium*. **(b)** To determine whether a resistance type is enriched in all genera, multivariate hypergeometric test (the lower, the more significant) is performed. The background colors are determined by the *p*-values measured by multivariate hypergeometric tests. **(c)** To determine among all genera containing *bac*A, which one is more *bac*A-enriched, a relative proportion calculation method is performed. 73.9% of all *bac*A sequences were found in *Flavobacterium*. **(d)** To determine whether a genus is enriched with ARGs, multivariate hypergeometric test is performed on each genus against all resistance types. *P*-values representing very significant ARG enrichment (*p*-value<1e-100) in four rows were highlighted in bold font, and so were the corresponding genus names (*Alteromonas, Pseudoalteromonas, Marinobacter*, and *Flavobacterium*). **(e)** To determine among all resistance types detected in genus *Flavobacterium*, which one is most enriched, the relative proportion calculation method is performed. The relative proportions of sequences of all 7 resistance types found in *Flavobacterium* and *bac*A sequences make up 41.58% of them, and it is highlighted by a red rectangle. Abbreviation: aac3ia, aac6ig Aminoglycoside Nacetyltransferase. acra, Resistance-nodulation-cell division transporter system. adeb, AdeB family multidrug efflux RND transporter permease. amrb, AmmeMemoRadiSam system protein B. ant3ia, Aminoglycoside O-nucleotidylyltransferase.aph33ib, streptomycin phosphotransferase. arna, Nucleoside-diphosphate-sugar epimerases. Baca, Undecaprenyl pyrophosphate phosphatase. bcra, Bacitracin transport ATP-binding gene. bl2a_nps, bl2b_tle, bl2c_bro, bl2d_oxa2, bl2e_y56: Class A beta-lactamase.catb1, catb2: Group B chloramphenicol acetyltransferase.

In above-mentioned taxonomy and biomarker analysis, many of the 11 ARG-containing genera were the members in the species co-occurrence network on genus level, indicating close connections among these genera. These genera had a clustering coefficient of 0.875, which was higher than the whole network clustering coefficient 0.744. Interestingly *Flavobacterium* (ARG-enriched) and *Polaribacter* (ARG-containing) were identified as hub nodes in the co-occurrence network. Top 4 ARG-enriched genera were all important biomarkers, with an average relative abundance above 0.1% in the 132 samples (**Supplementary Table S1**).

In the top 4 ARG-enriched genera, *Flavobacterium* was an important biomarker and hub node, it might have extensive interactions with other species, and the ARGs in *Flavobacterium* might protect it from antibiotics produced by other organisms in the same environment. Resistance type *bac*A was observed in several genera, but it drew our attention due to its enrichment in *Flavobacterium*, which was confirmed by both univariate and multivariate hypergeometric tests. We also found that 73.9% of all 66 *bac*A sequences were from *Flavobacterium* (**Fig. 3c**), and 41.58% of ARGs from *Flavobacterium* were *bac*A (**Fig. 3e**).

It has been shown that genus *Flavobacterium* plays an important role in community carbon cycling [31]. And the production of *bac*A shows undecaprenyl pyrophosphate (key component in cell wall biosynthesis) phosphatase activity and thus confers resistance to bacitracin that inhibits dephosphorylation [37]. With the metabolism production to develop the cell wall against the bacitracin, bacA shows the protective function as an ARG indirectly rather than inhibit the bacitracin itself. And as bacA gene was located on the chrome of *Flavobacterium*, which could encode protein effectively and was more stable than genes in plasmid [38]. Combing taxonomical analysis and ARG analysis, *bac*A might account for the role of *Flavobacterium* as a community hub and in carbon cycling, and previous genome analysis results showed that *bac*A indeed had been annotated in *Flavobacterium* [38].

### ARG impact on microbial community structure

In order to further analyze how ARGs affected the bacterial community, we constructed a phylogenic tree of 1,405 marine microbial genera (**Fig. 4a**) that we have identified (see **Supplementary File** for details), including 82 archaea and 1,323 bacteria. Based on the resulting phylogeny, we extracted 8 subtrees for the 11 ARG-enriched genera and their closest neighbors (**Fig. 4b**); in total 42 genera were included in the 8 subtrees. Within each subtree, pairwise *t*-tests were used to compare the relative abundances between the two species of each possible pairs across all 132 samples. We found that these ARG-enriched genera were all significantly more abundant than their ARG-less neighbors in the subtrees (*p*-value<0.01).

**Figure 4.**
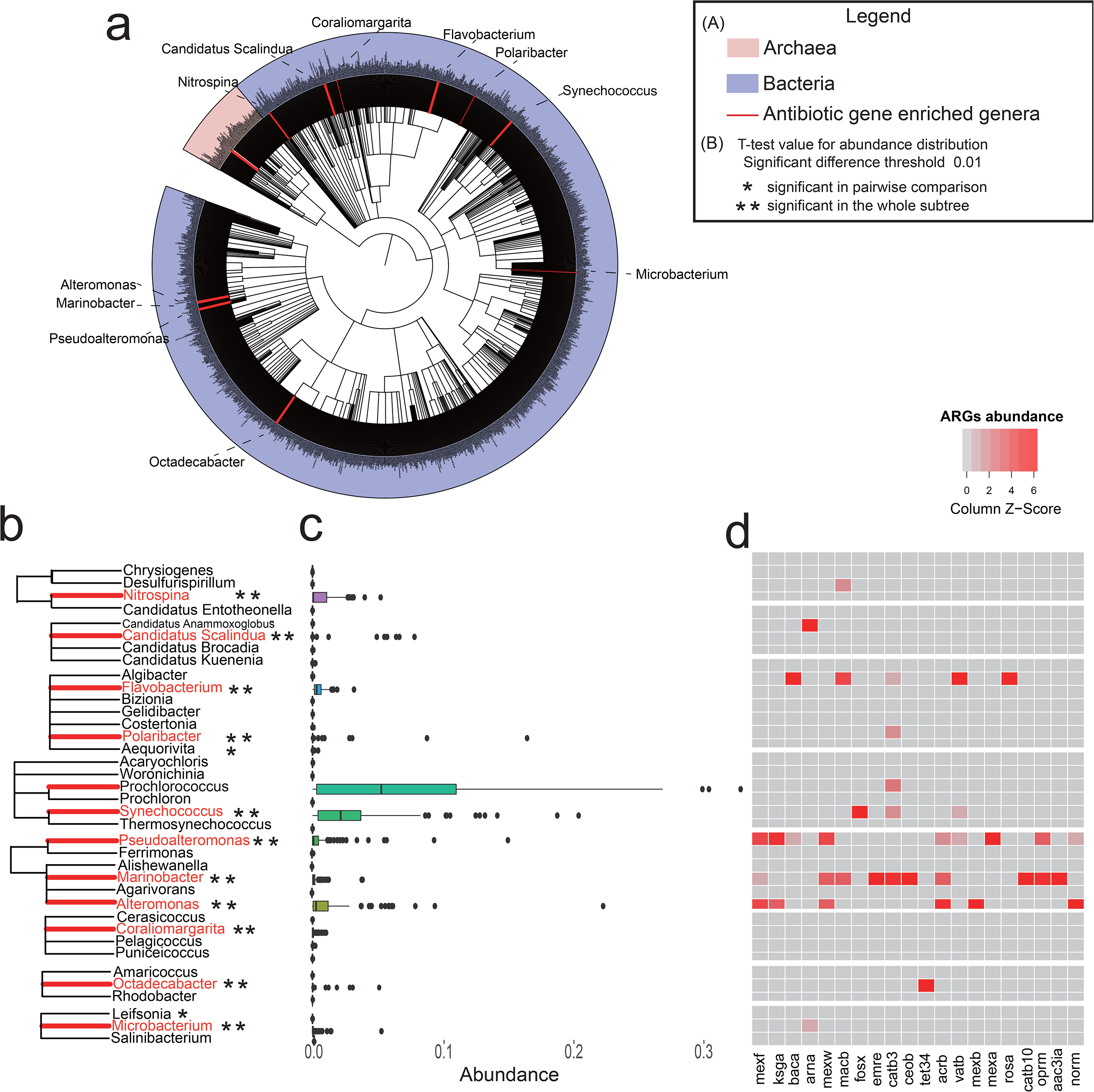
Phylogenetic analysis of ARG-enriched genera and their corresponding relative abundance and ARG enrichment patterns. **(a)** A phylogenetic tree of 1,405 detected marine genera, including archaea and bacteria. Branches colored red represent the phylogenetic locations of 11 ARG-enriched genera. **(b)** 8 subtrees containing the 11 ARG-enriched genera (highlighted by red lines) were selected from the phylogenetic tree, which in total contains 37 genera. These genera are enriched with ARGs compared with their closest phylogenetic neighbors (*) or all in the whole sub-tree (**). **(c)** Relative abundance of each of the 37 genera in (b) in 132 datasets (horizontally aligned). **(d)** A heatmap of the relative abundance distribution of several resistance types in the 37 genera in (b) (horizontally aligned). Horizontal axis represents the resistance types mapped to the genera in (b). Panels (b), (c) and (d) together indicate that genera enriched with ARGs are significantly more abundant in a microbial community, as well as compared with their phylogenetic neighbors in the microbial community.

More importantly, genera with close evolutionary relationship (i.e. neighbors in the subtrees) typically exist in similar environments [39]. However, on the 8 subtrees in **Fig. 4b**, the genera in the same subtree had a significant abundance difference in the marine bacterial communities (**Fig. 4c**). Combining the ARG distribution of the 37 genera, we found that genera with more ARGs had a higher abundance in the bacterial community (**Fig. 4d**). Therefore, our results indicated that ARG-enriched genera have a competitive advantage over ARG-less genera in the same environment.

In ocean environment, the ARGs could not only confer the antibiotics, but also had specific metabolic functions for ARG-enrichment genera [40], such as enzymatic synthesis, protein modification and metabolites degration to protect the bacteria from outside attack. For example, the ARG *bac*A enriched in *Flavobacterium* and take part in the cell wall development.

### Abundance of ARGs in Mediterranean samples implies a human factor

We next investigated if the abundances of ARGs in different samples could be (at least partially) influenced by human activities. Our hypothesis on how human activities could impact ARG contents and the community structure is illustrated in **Fig. 5a**. As we mentioned earlier, antibiotics used in Antimicrobial-producing industries, agriculture and House-hold waste may partially end up in the ocean through drainage and rainfall. Aquaculture, Antimicrobial-producing industries wasted water may directly Increase the amount of antibiotics into the ocean. And Antibiotics can be diluted easily in the open ocean [41], but not so in more closed water such as Mediterranean, especially when the latter is surrounded by human activities. The presence of antibiotics in the ocean may change the dynamic balance between naturally occurring antibiotics and ARGs [42], and will change the community structure by either killing some species that have no resistance to them [14], or promoting the exchange of ARGs among species [15, 16] that will also alter the community structure in the long term, or both. Consistent to our hypothesis, previous studies reported an increased anthropogenic impact on the antibiotic resistance profile in river estuary [43],[44].

In our study, we found that the average relative quantity (detailed normalization method in **Materials and Methods**) of ARGs detected in Mediterranean (the value is 7.18e-4) was noticeably higher than that in South Atlantic Ocean (the value is 2.13e-11). The reason behind might be that Mediterranean was enclosed water and near to the in-shore source of human-caused antibiotic content increase [45], while South Atlantic Ocean was more open and less impacted by human activities [46]. Alpha diversity analysis for species diversity of an environment also supported the potential effect of human-activity on in-shore ARGs: the average of both Shannon index and Simpson index are lower in Mediterranean than in South Atlantic Ocean (0.811 versus 0.906, and 0.333 versus 0.386 for the two indexes, respectively). As we have showed in **Fig. 4**, ARG-enriched bacteria could have competitive advantages over ARG-less species; this would be true especially when antibiotics are present (as illustrated in **Fig. 5c and 5d**). The difference indicated that environmental factors and human activities might be a key factor affecting ARG contents as well as microbial community structures [47].

**Figure 5.**
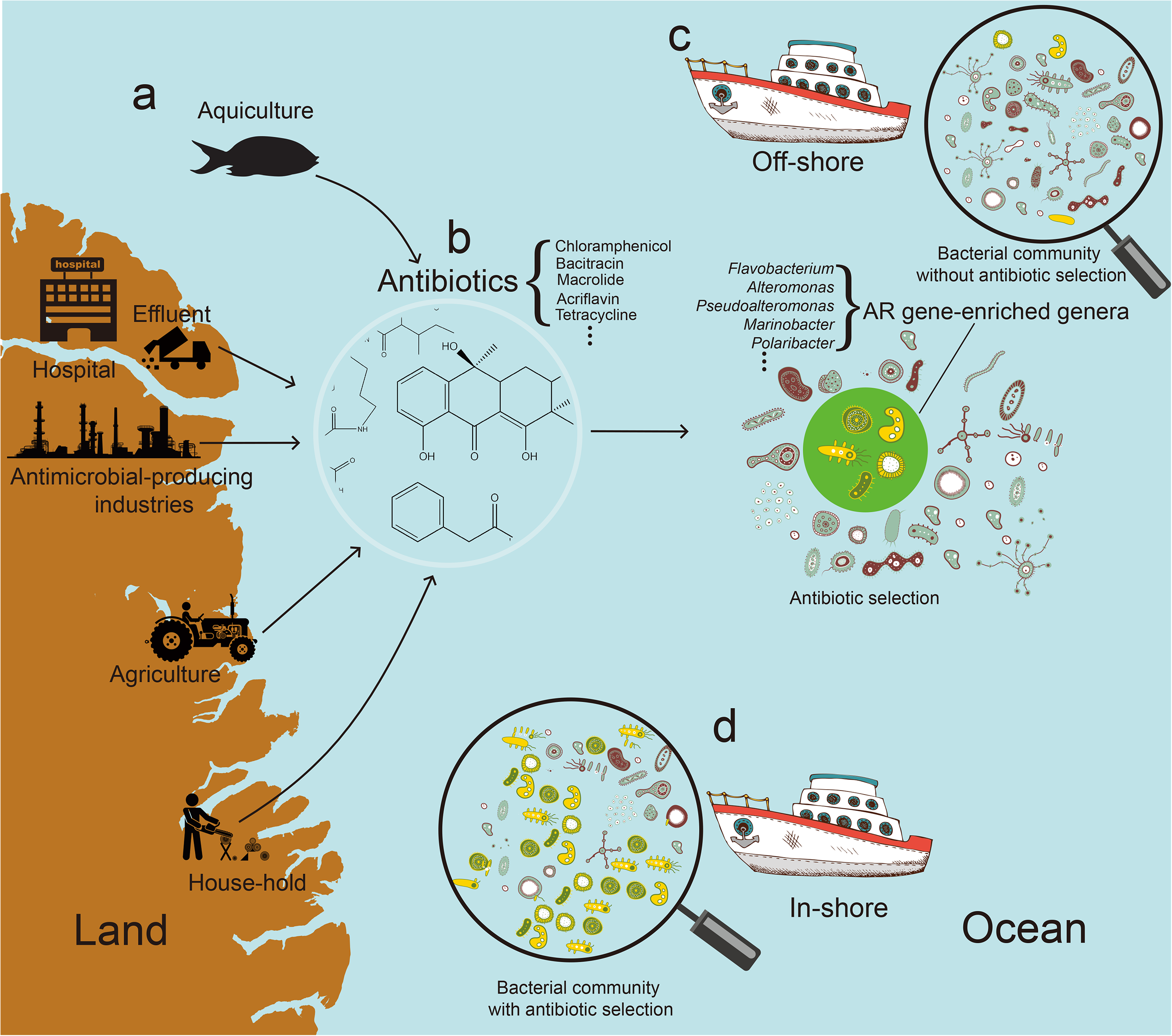
The hypothesis on possible involvement of human-activities in ARG influence on microbial community structures. **(a)** Possible antibiotic sources that are related with human activities. **(b)** ARGs might then become enriched in microbial communities under the selection pressure caused by antibiotics. **(c)** In an off-shore microbial community with little impact from antibiotics and human activities, the yellow colored genera in the green circle are not dominant. Genera shown here were identified as ARG-enriched by enrichment analysis (*Alteromonas, Pseudoalteromonas, Marinobacter*, and *Flavobacterium*, etc.). **(d)** An in-shore microbial community in which ARG-enriched genera (colored in yellow) become dominant.

## Conclusion

In this work, we reanalyzed the 132 metagenomic samples from the *Tara* Oceans project. Firstly, datasets grouped by different strategies have been compared, with results showing that water temperature, geographical locations and depth have exerted significant effects on the structure and functional profiles of the communities. Secondly, we have found biomarkers that were highly related with temperature (*Synechococcus* and *Prochlorococcus*, tending to live in warmer places), locations (*Planctomyces*, enriched in Atlantic Ocean), and depth (*Nitrospina* and *Alteromonas*, enriched in deeper layers). Thirdly, the analysis of species-species associations has revealed that the species co-occurrence patterns were heavily dependent on their environments. Finally, thousands of unique ARGs were identified, whose distribution patterns differ greatly by geographical locations and temperature. We found that ARG-enriched genera, such as *Alteromonas, Pseudoalteromonas, Marinobacter*, and *Flavobacterium*, were often more abundant than their ARG-less members in the same environment. More interestingly, an ARG against bacitracin (*bac*A), which was found in genus *Flavobacterium*, is pervasive in various environments, indicating that genera enriched with ARGs might possess an advantage over others in the competition for survival in the oceanic microbial communities.

Our study showed that deep mining of public marine metagenomic data could be useful for better understanding of the associations between community structures and functions of their key genes (e.g. ARGs). We believe that more profound associations and even causal relationships or patterns could be discovered by appropriate utilization of such resources and equally important by applying advanced data-mining techniques. In light of this, such integration of biotechnology (metagenomics) and information technology (data mining) would still need more high-quality multi-scale omics data. For example, such approaches might help us for better understanding of the process and significance on how human activities might affect ARGs, and subsequently affect the bacterial communities.

## Abbreviation

ARG: antibiotic resistance genes
WGS: whole genome sequence
*bac*A: Bacitracin Transport ATP-binding Gene
KEGG: Kyoto Encyclopedia of Genes and Genomes
OTU: Operational Taxonomic Unit
ARDB: Antibiotic Resistance Genes Database
SRF: Surface Water Layer
DCM/MIX: Subsurface Epipelagic Mixed Layer
MES: Mesopelagic Zone
*mex*F, *mex*B, *ceo*B: Multidrug Resistance Efflux Pump
*acr*B: Acriflavin Resistance
*ksg*A: Kasugamycin Resistance
EBI: The European Bioinformatics Institute
SPO: South Pacific Ocean
NPO: North Pacific Ocean
RS: Red Sea
MS: Mediterranean
SIO: South Indian Ocean
NIO: North Indian Ocean
NAO: North Atlantic Ocean
SAO: South Atlantic Ocean
PCC: Pearson Correlation Coefficient

## Declarations

### Availability of data and materials

A total of 132 metagenomic samples of Tara Oceans Project ERP001736 hosted on EBI Metagenomics were downloaded (https://www.ebi.ac.uk/metagenomics/projects/ERP001736)

### Author Contributions

Houjin Zhang and Kang Ning designed this study; Shiguang Hao, Chaoyun Chen and Pengshuo Yang collected and organized datasets; Shiguang Hao, Pengshuo Yang, Maozhen Han, Junjie Xu and Shaojun Yu analyzed the data; Shiguang Hao, Pengshuo Yang, Maozhen Han and Shaojun Yu interpreted the results; Shiguang Hao, Pengshuo Yang, Wei-Hua Chen, Houjin Zhang and Kang Ning wrote the initial draft of the manuscript; Shiguang Hao, Pengshuo Yang, Maozhen Han, Wei-Hua Chen, Houjin Zhang and Kang Ning revised the manuscript; all authors have read and approved the manuscript.

### Consent for publication

Not applicable

### Ethical Approval and Consent to participate

Not applicable

## Materials and Methods

### Datasets and categorizing strategies

A total of 132 metagenomic samples of *Tara* Oceans Project ERP001736 hosted on EBI Metagenomics were downloaded (https://www.ebi.ac.uk/metagenomics/projects/ERP001736) (**Supplementary Table S9**). These datasets were processed using the EBI Metagenomics pipeline (https://www.ebi.ac.uk/metagenomics/pipelines/2.0) prior to our downloading. The physical/chemical information was retrieved from the project site on EBI Metagenomics, and the geographical information was obtained from the supplementary file of ref. [10].

To analyze the correlations of environmental factors and taxonomical and functional profiles, we manually categorized the 132 samples into different groups according to their environmental attributes (**Supplementary Table S9**, **Supplementary Fig. S8**). We used 5 different attributes, namely depth (L, H), temperature (L1, L2, H1, H2), chlorophyll concentration (L1, L2, H1, H2), oxygen concentration (L1, L2, H1, H2), and geographical locations to group the 132 samples into distinct subgroups. For each attribute, the number of subgroups was indicated in the parenthesis; for the geographical location, the 132 samples were first divided into two groups and then in total eight sub-groups: the first group included samples from South Pacific Ocean (SPO), North Pacific Ocean (NPO), Red Sea (RS), and Mediterranean (MS), while the second group included samples from South Indian Ocean (SIO), North Indian Ocean (NIO), North Atlantic Ocean (NAO), and South Atlantic Ocean (SAO); datasets without such information were removed from subsequent analysis. Each resulting group contains similar number of datasets, with one exception that only five datasets are in the group of shallow area with a low oxygen concentration due to high temperature. The detailed categorizing criteria and results are shown in **Supplementary Table S10-S13**.

### Taxonomical and functional profiling of metagenomic datasets

#### Analysis of taxonomical and functional profiles

For each dataset, 16S rDNA sequence reads were extracted from processed reads using Parallel-Meta v3.2.1 [48]. The files containing the 16S rDNA sequences (in fasta format) were used as input data and submitted to Parallel-Meta. By aligning non-chimeric reads to the Greengenes database (v13_5) [49], the OTUs were obtained based on a sequence similarity cut-off of 97%. Sensitive alignment mode and Fwd & Rev pair-end sequence orientation were used. Other parameters were kept default. Based on the taxonomical structures and relative abundance of communities, functional annotations at phylum, genus, and Operational Taxonomic Unit (OTU) levels were analyzed according to Kyoto Encyclopedia of Genes and Genomes (KEGG) [28]. Alpha diversity statistical methods including Shannon index, Simpson index were used for 132 samples.

#### Construction of co-occurrence network on species level

To characterize the microbial communities comprehensively, network analysis was performed on phylum, genus, and OTU levels. As relative abundances of species were calculated by Parallel-Meta, only those with abundances above 0.01% were kept for network construction. Species co-occurrence matrix was generated using in-house C++ scripts, calculated by making the quantitative comparison between species using the Pearson Correlation Coefficient (PCC) for each pair of bacteria. The PCC threshold at different levels was set to ±0.10, ±0.10, and ±0.50, respectively. For choosing reasonable method to calculate the species co-occurrence correlation, the alpha diversity in taxonomy analysis and abundance distribution on OTU level were considered [50]. With average Simpson index of 0.99 and more than 50% sparse after filtering to remove very rare OTUs, Pearson correlation was reasonable for bacteria data without time series. A species co-occurrence matrix including all qualified pairwise PCC was generated and imported to Cytoscape v3.4.0 for further analysis [51]. MCODE algorithm was used as a clustering method for network analysis [52]. When degree was >2 and node score was >0.2, the node was clustered. The largest depth for clustering was 100. Other parameters were set as defaults.

### Metagenomic assembly and prediction of antibiotic resistance genes

The processed reads were assembled and processed by using DESMAN [53], with nextflow pipeline to perform the reads assembly and contig binning. With a collection of 37 genes from bacteria and archaea to identify contig bins, the species distribution in 132 samples could be calculated.

A protein reference file was downloaded from Antibiotic Resistance Genes Database (ARDB, http://ardb.cbcb.umd.edu/) [33]. Entries with 100% identical sequences were merged, and three nucleotide sequences that are not indexed in ARDB website were removed. After being cleaned up, the reference contained 2,893 translated sequences of ARGs. Blastx searching was performed with an e-value threshold of 1e-10. A query sequence was annotated as an ARG if the first high-score pair (HSP) of its top hit showed a percent identity ≥60% and a query coverage ≥70%.

The number of unique ARGs detected in each dataset was normalized by the number of reads (representing the data size of the sample) and the number of OTUs (representing the complexity of the sample) in that dataset. 

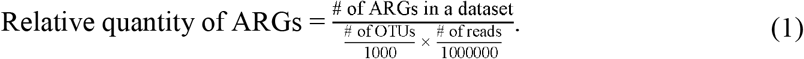

The number of resistance types in each dataset was normalized according to equation. 

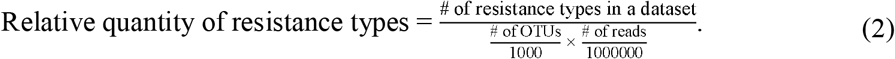

### Antibiotic resistance gene enrichment in marine microbial genera

Twenty-four genera were selected for this analysis, each having an average abundance above 0.1% among samples. Of these genera, “HTCC” and “SargSea-WGS” were abandoned due to their ambiguous names. Records related to the remaining 22 genera in the NCBI nr database (retrieved on 24th Nov, 2016) were extracted and filtered, and 2,919,490 unique accessions were obtained. BLASTp searching against the NCBI nr database was performed and restricted among these accessions. The e-value threshold was set to 1e-10. For each query sequence, the organism name of its top hit subject sequence was assigned to it, if the percent identity is ≥40%. In cases where the subject sequence has multiple organism names on record, the first one was selected.

The enrichment analysis was performed as below. 1) To determine whether a resistance type is enriched in a genus, we performed univariate hypergeometric test on each resistance type against each genus using Scipy module in Python (http://www.scipy.org/). 2) To determine whether a resistance type is enriched in all genera (*p*-value<0.01), we performed multivariate hypergeometric test on each resistance type against all genera using R package BiasedUrn v1.05 (https://cran.r-project.org/web/packages/BiasedUrn/). Central multivariate hypergeometric distribution model was used in the calculation of *p*-values. 3) To determine whether a genus is enriched with ARGs of all resistance types when compared with other genera, we performed multivariate hypergeometric test on each genus against all resistance types using BiasedUrn based on central multivariate hypergeometric distribution model. 4) To determine among all genera containing *bac*A, which one is more *bac*A-enriched, we introduced a relative proportion calculation method: The quantity of *bac*A sequences in each *bac*A-containing genus was counted, and the results were normalized (dividing the number of *bac*A sequences of this genus, by the total number of *bac*A sequences for all genera) and illustrated. 5) To determine among all resistance types enriched in genus *Flavobacterium*, we again used the relative proportion calculation method in 4). The quantity of all ARGs from *Flavobacterium* were counted, and the results were normalized (dividing the number of *bac*A sequences of *Flavobacterium*, by the total number of ARG sequences of *Flavobacterium*) and illustrated.

In order to uncover the association of human activities, ARGs, and microbial communities, a phylogenetic tree including 1,405 detected marine genera in 132 samples was constructed at genus level, then the abundance and ARG distribution of ARG-enriched genus and their neighbors in the same subtree were compared. There are in total 1,664 genera identified by Parallel-Meta; after removing genera with multiple taxonomy IDs from the NCBI taxonomy database [54] and manually adding some genera with conflicting names in Parallel-Meta and NCBI taxonomy database, we obtained 1,405 genera with a validated NCBI taxonomy ID (detailed genera and taxa ID see **Supplementary File**). PhyloT (http://phylot.biobyte.de/) was used to map the 1,405 taxonomy IDs to the NCBI common tree (https://www.ncbi.nlm.nih.gov/Taxonomy/CommonTree/wwwcmt.cgi), and subsequently the results were visualized and modified by an online tool iTOL [55] and Evolview [56]. 9 subtrees containing the ARG-enriched genera (42 genera) were selected. Boxplots that show the abundance distribution of the 42 genera across the 132 datasets were plotted next to the subtrees. A heatmap of ARG count distribution in all the 42 genera was plotted and the values in each column were normalized using *z*-score.

